# β-Lactamase-Responsive Probe for Rapid and Non-invasive Detection of Tuberculosis through Induced Breath Analysis

**DOI:** 10.64898/2026.03.03.709436

**Authors:** Guochao Wu, Dongfang Zhou, Han Jin

**Affiliations:** Institute of Nano Biomedicine and Engineering & National Key Laboratory of Advanced Micro and Nano Manufacture Technology, School of Automation and Intelligent Sensing, Shanghai Jiao Tong University, Shanghai 200240, P. R. China; National Engineering Research Center for Nanotechnology, Shanghai 200241, P. R. China; NMPA Key Laboratory for Research and Evaluation of Drug Metabolism & Guangdong Provincial Key Laboratory of New Drug Screening & Guangdong-Hongkong-Macao Joint Laboratory for New Drug Screening, School of Pharmaceutical Sciences, Southern Medical University, Guangzhou 510515, PR China; Department of Pharmacy, The Seventh Affiliated Hospital, Southern Medical University, Foshan 528244, China

**Keywords:** pulmonary tuberculosis (PTB), cefapirin sodium (CS), hydrogen sulfide (H_2_S), β-lactamase (Blac), Gas sensor, Induced breath diagnosis

## Abstract

Pulmonary tuberculosis (PTB) remains a major global public health challenge. Existing diagnostic approaches are generally limited by suboptimal sensitivity, prolonged turnaround times, high costs, and reliance on sputum samples. These constraints hinder large-scale implementation in resource-limited settings and substantially impede early screening and timely intervention. Targeting the characteristic expression of β-lactamase in Mycobacterium tuberculosis, this study employed systematic screening of a molecular compound library and identified cefapirin sodium as a specific molecular probe. Upon selective hydrolysis by β-lactamase, the probe releases hydrogen sulfide (H_2_S), which can be detected using a highly sensitive H_2_S sensing system, thereby enabling rapid and noninvasive identification of the pathogen. Gas chromatography-mass spectrometry (GC-MS) and colorimetric assays were used to validate the specificity of the enzymatic reaction and to confirm the accuracy of the generated product, collectively demonstrating the feasibility of the proposed detection platform. This method offers several advantages, including noninvasiveness, rapid response, and low cost, making it particularly suitable for application in resource-constrained regions. It provides a promising new strategy for early, point-of-care, and large-scale screening of pulmonary tuberculosis, with important implications for improving TB control efforts in high-burden settings.

## 1. Introduction

Pulmonary tuberculosis (PTB) remains a major global public health challenge and one of the leading causes of mortality from infectious diseases.^1-3^ According to the World Health Organization, millions of new tuberculosis cases are reported annually, with a substantial proportion occurring in high-burden countries such as India, Indonesia, and South Africa.^4-6^ Despite continuous advances in chemotherapy and molecular diagnostics, delayed case detection continues to sustain community transmission. PTB is caused by *Mycobacterium tuberculosis*, an intracellular pathogen that invades and survives within host macrophages in the lung.^7^ Notably, *M. tuberculosis* expresses β-lactamase (Blac), an enzyme capable of hydrolyzing β-lactam antibiotics by cleaving the β-lactam ring, thereby reducing drug efficacy and contributing to intrinsic resistance.^8^ Together with inappropriate treatment and transmission of resistant strains, this mechanism facilitates the emergence and spread of multidrug-resistant tuberculosis (MDR-TB), further complicating disease control.^9, 10^

In many resource-limited, high-burden settings, sputum smear microscopy remains widely used as a primary diagnostic tool.^11^ However, this method suffers from limited sensitivity, particularly in paucibacillary cases, leading to false-negative results.^12^ In addition, conventional culture-based confirmation--although more sensitive--may require several weeks, sometimes up to eight weeks, to yield definitive results.^13, 14^ The introduction of rapid molecular assays such as Xpert MTB/RIF has markedly improved diagnostic efficiency by enabling detection of the *M. tuberculosis* complex and rifampicin resistance-associated mutations within approximately two hours.^15-17^ Nevertheless, this platform remains largely dependent on sputum specimens and requires relatively costly instrumentation and stable infrastructure, posing challenges for sustained implementation in remote and resource-constrained regions. Moreover, molecular assays are not readily adaptable for real-time, continuous, or non-invasive population screening. In high-burden areas, additional barriers--including difficulty in sputum collection, suboptimal patient compliance, and delays in healthcare access--further hinder early case identification. Timely diagnosis of PTB is therefore essential for interrupting transmission chains and improving clinical outcomes.

Against this backdrop, non-invasive diagnostic strategies based on exhaled breath analysis have emerged as promising alternatives.^18, 19^ Pulmonary infection and host immune responses can perturb metabolic pathways, resulting in the production of specific volatile organic compounds (VOCs) and gaseous signaling molecules that are detectable in exhaled breath.^20, 21^ Breath-based diagnostics offer several advantages, including non-invasiveness, rapid sampling, minimal biosafety risk, and potential applicability to point-of-care testing and large-scale screening.^22, 23^ Accordingly, the development of rapid, highly sensitive, and cost-effective breath-based detection technologies represents a strategically important direction for achieving early PTB diagnosis and strengthening global tuberculosis control efforts.

To address the aforementioned challenges, we developed a novel detection strategy for *M. tuberculosis* based on its specific expression of Blac. Exploiting the enzymatic ability of

Blac to hydrolyze the β-lactam ring, we screened a molecular compound library to identify suitable substrates and found cefapirin sodium (CS, a clinically used antibiotic containing a β-lactam ring structure) to be an ideal molecular probe. Upon enzymatic hydrolysis by Blac through a cascade reaction process, this compound undergoes structural transformation leading to the release of hydrogen sulfide (H□S), a biologically relevant gaseous signaling molecule. The generated H□S can then be detected using a specific hydrogen sulfide gas sensor, thereby enabling rapid detection of *M. tuberculosis* (Scheme 1). If successfully translated into clinical practice, this strategy has the potential to achieve rapid detection, high analytical sensitivity, and simultaneous identification of drug-resistance-related enzymatic activity. Such an approach could substantially shorten diagnostic delays, interrupt transmission chains, and improve tuberculosis control outcomes in high-burden settings.

**Scheme 1.**
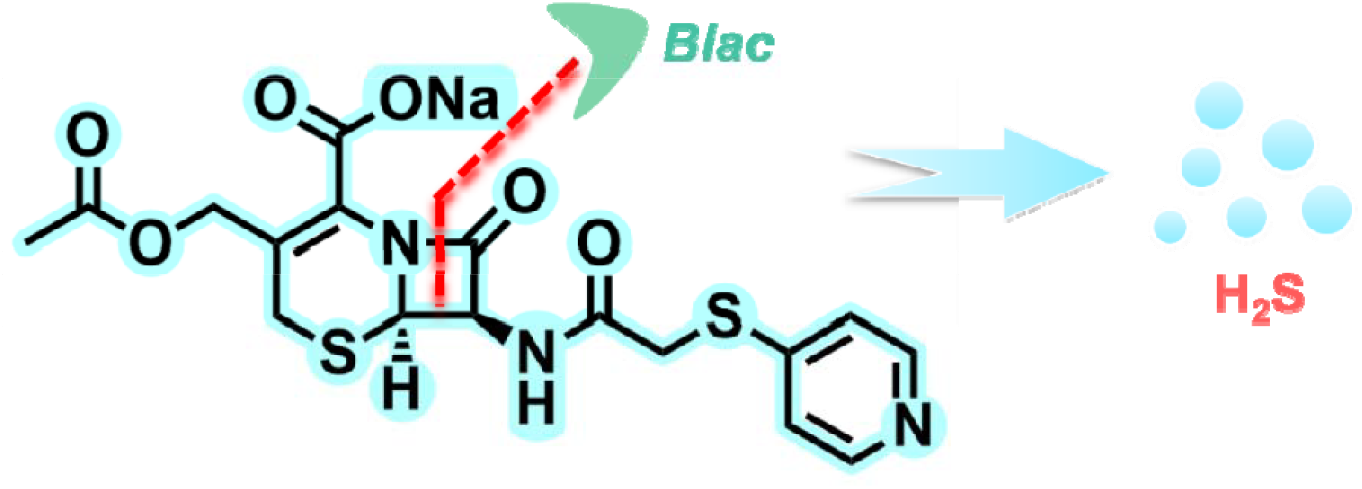
Schematic illustration of the principle of cefapirin sodium as a molecular probe for early detection of pulmonary tuberculosis.

## 2. Results and discussion

To validate the core feasibility of the proposed early detection strategy for PTB, an in vitro reaction system was first established to investigate the interaction between CS and the *M. tuberculosis*--specific Blac. Three parallel groups were designed: (i) a negative control containing Blac alone, (ii) a positive control containing CS alone, and (iii) an experimental group in which Blac was co-incubated with CS. All reaction mixtures were incubated in a thermostatic water bath at 37.5 °C for 1 h. Subsequently, volatile components in the headspace were qualitatively analyzed using gas chromatography-mass spectrometry (GC-MS). The corresponding results are presented in Figure 1A-C. A distinct characteristic H□S signal peak was observed exclusively in the extracted ion chromatogram of the experimental group, whereas no such peak was detected in either the negative or positive control groups. These findings indicate that H□S release is dependent on the specific enzymatic reaction between Blac and CS. Further mass spectral comparison demonstrated that the characteristic peak detected in the experimental group exhibited a high degree of concordance with the reference mass spectrum of H_2_S in the NIST14 spectral library (Figure 1D, E).

**Figure 1.**
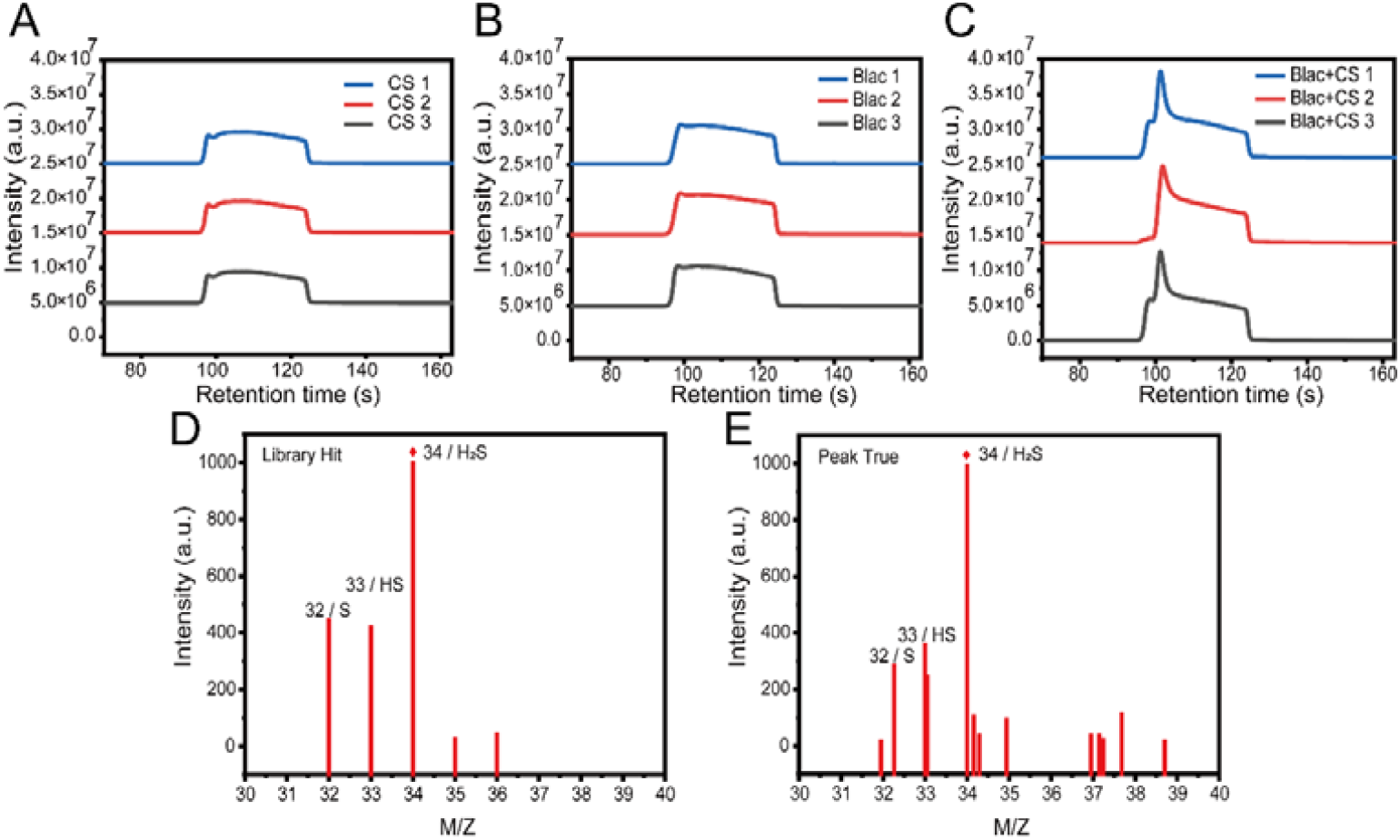
Enzymatic assay of H_2_S release from the drug under Blac catalysis. (A, B): Neither CS nor Blac alone produces H_2_S after incubation in a water bath. CS and Blac exhibit a certain degree of stability. (C) : Co-incubation of CS and Blac results in a significant H_2_S signal. (D, E): Mass spectra of H_2_S obtained from the standard spectral library (D) and sample extraction (E).

Subsequently, H□S signal peak obtained from the enzymatic reaction system by GC-MS analysis was compared with the calibration curve generated using standard H□S gas (Figure 2A, B). The results demonstrated that the characteristic peaks of the experimental group were highly consistent with those of the standard in both retention time and relative abundance. These findings further confirm that CS can specifically generate H_2_S upon catalytic activation by Blac.

**Figure 2.**
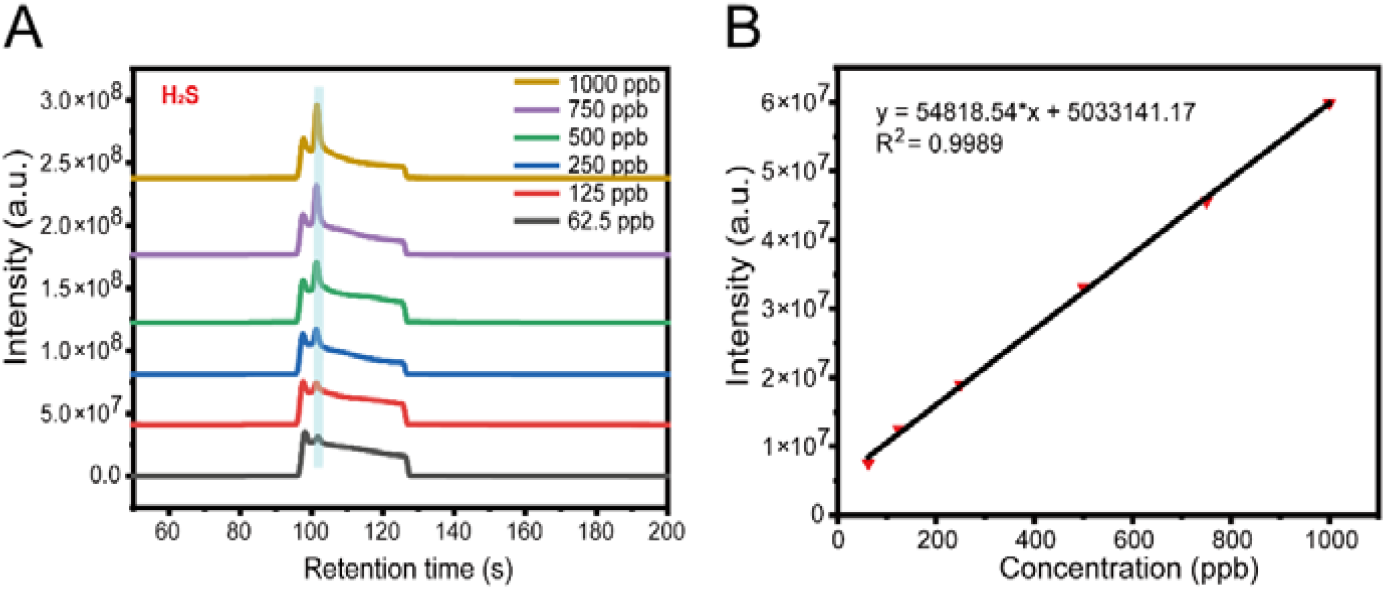
Calibration curve for H_2_S concentration. (A) Detection results of H_2_S gas by headspace solid-phase microextraction coupled with gas chromatography-mass spectrometry (HS-SPME&GC-MS), with a concentration range of 62.5-1000 ppb. (B) Fitted curve of H_2_S concentration versus peak intensity.

To enable more direct visualization of H□S production, the standard methylene blue assay was employed for colorimetric detection.^24^ In this system, H□S undergoes a specific chromogenic reaction, resulting in a color change of the solution from colorless to blue and the emergence of a characteristic UV-vis absorption peak at 665 nm. As shown in Figure 3A, the co-incubation group containing Blac and CS exhibited a pronounced absorption peak at 665 nm, with both peak shape and position highly consistent with those observed in the H□S standard solution group. In contrast, no significant absorption signal was detected in the control groups treated with CS alone or Blac alone, indicating that H□S generation depends on the specific enzymatic interaction between the enzyme and its substrate. Moreover, within a defined concentration range, a progressive increase in the UV-vis absorption intensity corresponding to the chromogenic reaction was observed with increasing concentrations of the substrate CS (Figure 3B), demonstrating a clear concentration-dependent enhancement. This finding further suggests a close correlation between reaction intensity and substrate concentration.

**Figure 3.**
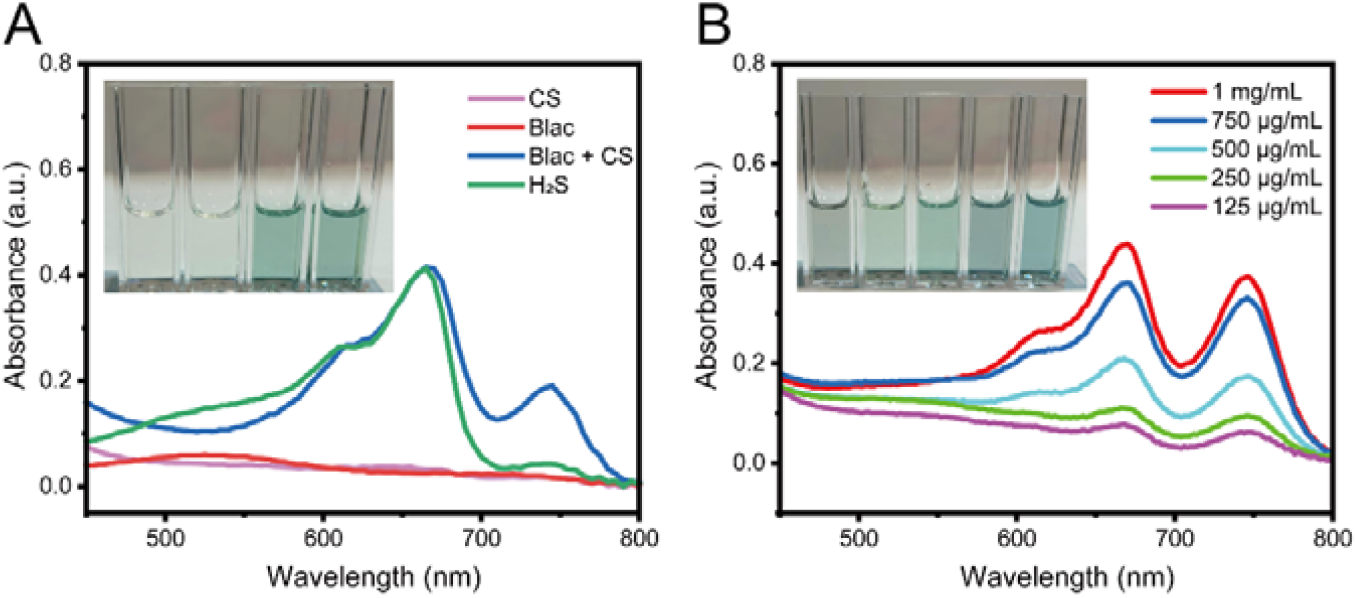
Determination of H_2_S production using the standard methylene blue assay. (A) UV-vis spectra with characteristic absorbance at 665 nm and corresponding photographs (inset) of CS, Blac, Blac + CS, and H_2_S solutions after addition of methylene blue. (B) Concentration-dependent UV-vis absorbance spectra and corresponding photographs (inset) of Blac incubated with varying concentrations of CS (1 mg/mL, 750 µg/mL, 500 µg/mL, 250 µg/mL, and 125 µg/mL) following methylene blue addition.

Collectively, these results corroborate from multiple perspectives that the specific enzymatic reaction between Blac and CS can stably generate H_2_S. This provides robust experimental support for the subsequent development of a noninvasive breath-based diagnostic platform leveraging this reaction, with the aim of enabling rapid and early detection of PTB.

## 3. Conclusions

In summary, to address the challenges associated with early PTB detection in economically underdeveloped regions, this study was designed to meet the core demands of primary healthcare settings by exploring a molecular probe-based strategy for early PTB diagnosis. Through systematic screening of a molecular compound library, CS was successfully identified as an ideal probe with specificity toward *M. tuberculosis*. CS undergoes an efficient enzymatic reaction with Blac specifically expressed by *M. tuberculosis*, resulting in selective cleavage of its β-lactam ring and subsequent release of H□S. By integrating a dedicated H_2_S gas sensor, changes in H□S levels in exhaled breath can be monitored to enable noninvasive and specific detection of PTB infection. To date, preliminary proof-of-concept validation and feasibility assessments have been completed, yielding stage-specific results that substantiate the specificity of CS as a molecular probe.

To further advance the clinical translation of this technology, future studies will focus on two major directions. First, comprehensive in vitro microbiological assays and in vivo animal model experiments will be conducted to systematically evaluate detection sensitivity, specificity, stability, and operational parameters, thereby defining the applicable scope and optimal testing conditions. Second, parallel efforts will be devoted to the optimization and development of a high-performance H_2_S gas sensor, with emphasis on improving response time, limit of detection, portability, and resistance to environmental interference, ultimately promoting device miniaturization and system integration. Through multidimensional experimental validation and iterative technological refinement, this platform is expected to progress from laboratory development to practical implementation in primary clinical settings, providing critical technical support for reducing missed diagnoses and improving the efficiency of early intervention in PTB.

## CRediT authorship contribution statement

**Guochao Wu**: Conceptualization, Data curation, Formal analysis, Investigation, Methodology, Validation, Visualization, Writing - original draft. **Dongfang Zhou:** Conceptualization, Formal analysis, Resources, Methodology, Writing - review & editing. **Han Jin:** Conceptualization, Formal analysis, Funding acquisition, Investigation, Methodology, Project administration, Resources, Supervision, Validation, Visualization, Writing - review & editing.

## Data availability

Data will be made available on request.

